# Making high-dimensional molecular distribution functions tractable through Belief Propagation on Factor Graphs

**DOI:** 10.1101/2021.06.28.450193

**Authors:** Zachary Smith, Pratyush Tiwary

**Affiliations:** Biophysics Program and Institute for Physical Science and Technology, University of Maryland, College Park 20742, USA; Department of Chemistry and Biochemistry and Institute for Physical Science and Technology, University of Maryland, College Park 20742, USA

## Abstract

Molecular dynamics (MD) simulations provide a wealth of high-dimensional data at all-atom and femtosecond resolution but deciphering mechanistic information from this data is an ongoing challenge in physical chemistry and biophysics. Theoretically speaking, joint probabilities of the equilibrium distribution contain all thermodynamic information, but they prove increasingly difficult to compute and interpret as the dimensionality increases. Here, inspired by tools in probabilistic graphical modeling, we develop a factor graph trained through belief propagation that helps factorize the joint probability into an approximate tractable form that can be easily visualized and used. We validate the study through the analysis of the conformational dynamics of two small peptides with 5 and 9 residues. Our validations include testing the conditional dependency predictions through an intervention scheme inspired by Judea Pearl. Secondly we directly use the belief propagation based approximate probability distribution as a high-dimensional static bias for enhanced sampling, where we achieve spontaneous back-and-forth motion between metastable states that is up to 350 times faster than unbiased MD. We believe this work opens up useful ways to thinking about and dealing with high-dimensional molecular simulations.

## I INTRODUCTION

Understanding the relationships between the many different degrees of freedom constituting a generic high-dimensional molecular system is a problem of longstanding theoretical and practical interest.^1–3^ For a system with *k* ≫ 1 degrees of freedom, in principle one could consider the full *k*–body probability distribution *P*, which would account for any and all correlations between the *k* different components. However, a full *k*–body probability distribution is not always necessary as can be intuited with the popularity of the Ising model, where nearest-neighbor interactions can capture arbitrarily long-range communication and correlations, and the full joint probability is not needed to model these relationships. Furthermore, fitting such a *k*–dimensional *P* to data would involve *k*–dimensional histograms which becomes impractical as *k* > 2 ~ 3. In fact, learning tractable probabilistic models for high dimensional data is a problem that extends far beyond molecular sciences and pervades numerous aspects of modern day life and data science.^4^ Understanding the dependencies in data is thus the first step toward understanding the processes that generated the data, and for further simulating these processes efficiently through advanced sampling methods for instance.^5^

In this work our specific interest is in data coming from Molecular dynamics (MD) simulations, a tool very commonly used to study the dynamics of chemical systems at all-atom spatial resolution and femtosecond temporal resolution. This leads to a deluge of high-dimensional data which can be hard to make sense of. Specifically for biomolecules, here we would like to characterize which residues are fluctuating in a mutually correlated manner, and use this information to better understand the governing biophysical processes. Furthermore, due to the high temporal resolution of MD, timescales associated with practical problems such as protein folding and drug unbinding are inaccessible even using most modern high power computing resources. Thus we are also interested in using these tractable probability distributions governing the high-dimensional MD to enhance the simulation itself.^6^ To do so we propose a framework using methods from probabilistic graphical models^4^ to construct a factorization of input degrees of freedom. Among other works in this area, our work is inspired by Ref. 7–8. Given our eventual interest in sampling slow, representative degrees of freedom, we take the liberty to call these degrees of freedom as order parameters (OPs). We chose to use probabilistic graphical models taking inspiration from their successes in a variety of fields ranging from identifying disaster victims^9^ to decoding messages^10^ where the relationships between unknowns are leveraged to perform an inference task with levels of efficiency that wouldn’t be possible otherwise.

The key ideas, detailed in Sec. II, can be summarized as follows. We start with high-dimensional data with complicated and *a priori* unknown dependencies between the various OPs, with the ultimate objective of approximating this intractable high-dimensional probability distribution with a factorized, tractable probability comprising up to triplet terms. For this, we first assess the conditional dependencies between the OPs using Markov random fields (MRF)^11^ detailed in section II A. MRFs are graphical models where non-adjacent nodes are conditionally independent. We use graphical lasso,^12^ an estimator for the inverse covariance matrix to determine the conditional dependence relationships used in the MRF. In Sec. II B we describe how these conditional dependency relationships are converted into a bipartite graph known as factor graph, where the two disjoint sets of vertices denote respectively OPs and the functions connecting them. These functions are then learnt using the belief propagation algorithm (Sec. II C).^8,13^ In order to assess the benefits of this factor graph-based approach we apply it to simulations of two well-studied small peptides^14–16^ Ala_3_ and Aib_9_ peptide that display rich conformational dynamics in spite of their small sizes (Fig. 1), and can be described using their *ϕ* and *ψ* dihedral angles. The factor graphs are validated using a range of additional simulations in Sec. III, which include (i) an intervention protocol in the sense of Pearl^17^ that directly confirms our predictions of conditional dependency between different degrees of freedom and (ii) a demonstration of how this knowledge can lead to up to three orders of magnitude enhanced sampling along all degrees of freedom, with hysteresis-free back-and-forth movement between different metastable states. We believe this work represents a new use of ideas from the field of probabilistic graphical models^4^ in the study of molecular simulations aiding with better interpretations and superior sampling.

**FIG. 1:**
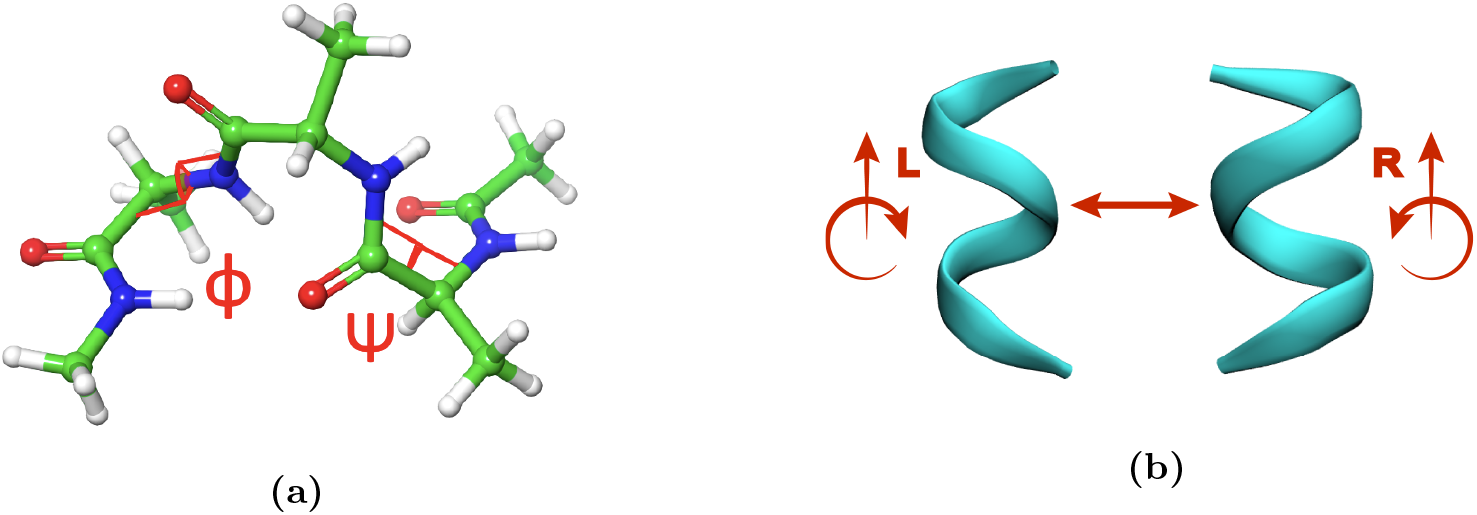
The two model systems simulated in this work capped Ala_3_ (a) and Aib_9_ peptide (b). Alanine peptide atoms are shown with an example of a *ϕ* and *ψ* angle overlayed in red. Aib_9_ peptide is shown as a ribbon to highlight its helical secondary structure.

## II METHODS

In this Section we describe the theory underlying our key ideas as summarized in the Introduction above, as well as the workflow for its implementation (Fig. 2).

**FIG. 2:**
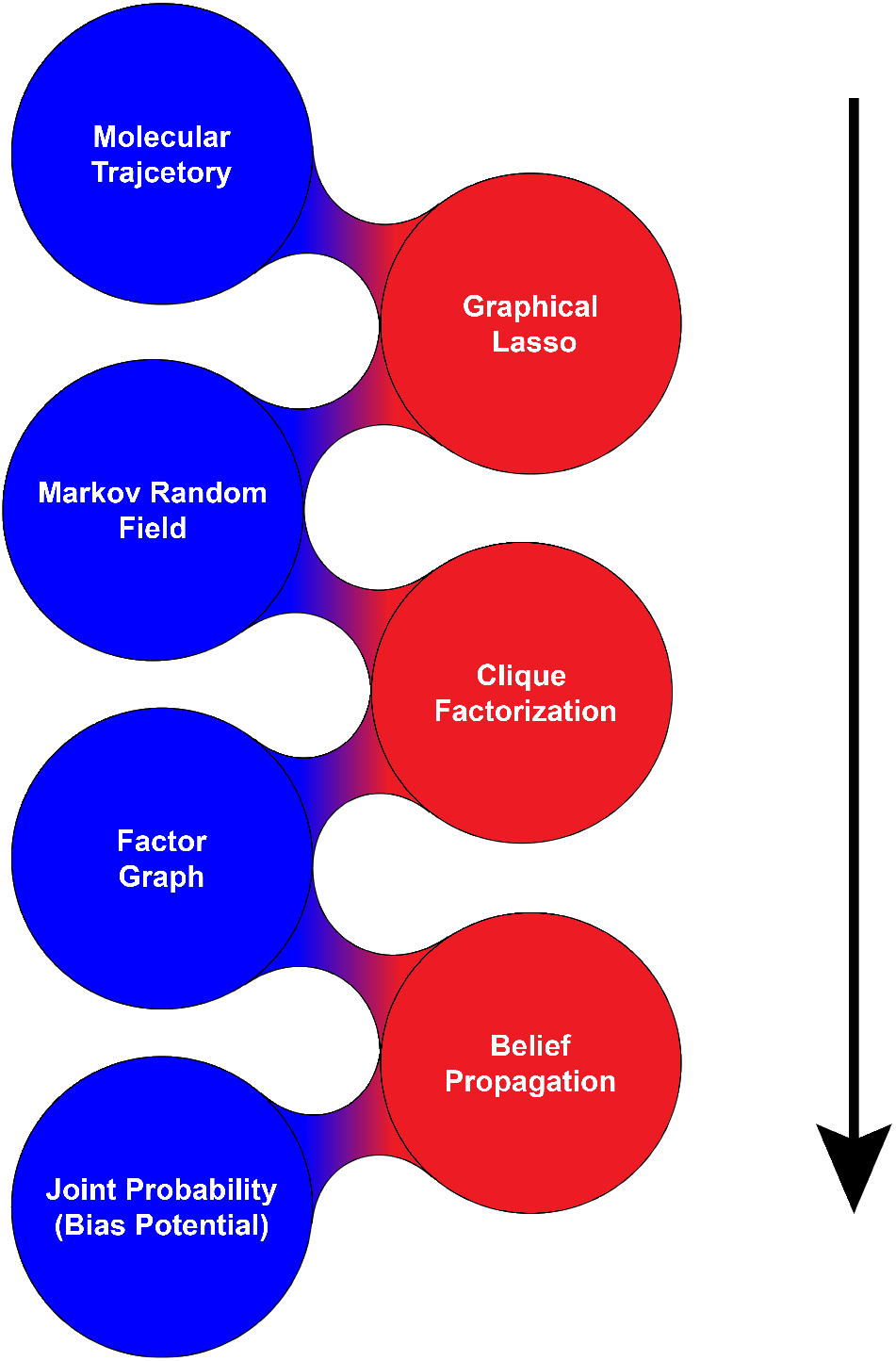
Schematic of the workflow done in this manuscript as a bipartite graph with objects shown in blue and algorithms shown in red. The workflow proceeds from top to bottom.

### II.A Markov Random Fields (MRF)

Our starting point is sampling of high-dimensional OPs arising from molecular dynamics (MD) simulations (see supplementary information (SI) for details) but the proposed procedure should be more generally applicable. In order to partition OPs in to simpler lower-dimensional factors, we first assess the relationships between the OPs themselves. Consider any 3 random variables A, B, and C. A particularly informative measure of the dependencies between such variables is conditional independence, denoted (*A* ⫫ *B*) *C*.^18^ This measure is used in the pairwise Markov property of Markov random fields (MRF), and denotes that the two variables *A* and *B* are statistically independent given knowledge of *C*. Mathematically this can be written as

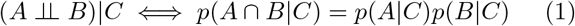

with *p* denoting respective probability densities. Here we call variables equivalently as nodes shifting to a graphical modeling perspective. Generalizing Eq. 1 to all possible nodes, the pairwise Markov property is satisfied if two nodes are conditionally independent given all other nodes as shown in Eq. 2, where *X_i_* and *X_j_* denote nodes and *X*_*N*\{*i,j*}_ denotes all other nodes.

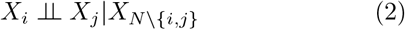

A MRF is an undirected graphical model such that all nodes without edges between them have the pairwise Markov property and those with edges do not have it.^4^ This can be interpreted as meaning that any correlation between connected nodes can not be “explained away” by a third node causing the correlation. The presence of the pairwise Markov property is estimated for all possible edges by estimating an inverse covariance matrix (Θ). All pairs of variables *X_i_*, *X_j_* that have a zero at the associated position in the inverse covariance matrix (Θ*_i,j_*) can be expected to have the pairwise Markov property.^4^ To efficiently calculate this inverse covariance matrix, we use the graphical lasso^12^ estimator shown in Eq. 3:

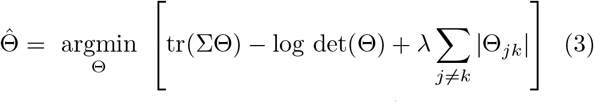

where Θ is the inverse covariance, 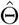 is the inverse co-variance returned by graphical lasso, Σ is the observed covariance, *λ* is a parameter that controls how strongly off-diagonal terms are penalized, tr is the matrix trace, and det is the matrix determinant. This *λ* parameter can be effectively used to control the sparsity of the resulting MRF as is shown later for our examples (Fig. 6). In practice, we tune this parameter to produce the most densely connected MRF that contains at most triplet interactions (in other words, the factor graph later described in Sec. II B should have clique size no larger than three).

### II.B Factor Graphs (FG)

Once we have a MRF, we can use this graph structure to partition the OPs into independent sets or factors that make up the joint distribution. It is important to note that these independent sets are not necessarily disjoint, and the same variable could be a part of different factors as we show for practical examples in Sec. III. The structure of these factors can be obtained from the cliques, or sets of fully connected nodes, of the MRF. Each maximal clique, or clique that is not part of a larger clique, contains the variables in one factor of the joint probability. This relationship is illustrated through a schematic in Fig. 3a. It is important to note that at this point we know the variable sets used for each factor but not their functional form. More technically, this is defined in Eq. 4 where *C*(*G*) is the set of all maximal cliques on a graph *G* and *f_c_*(*X_c_*) is each factor as a function of the variables contained in the associated clique:

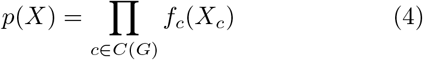

**FIG. 3:**
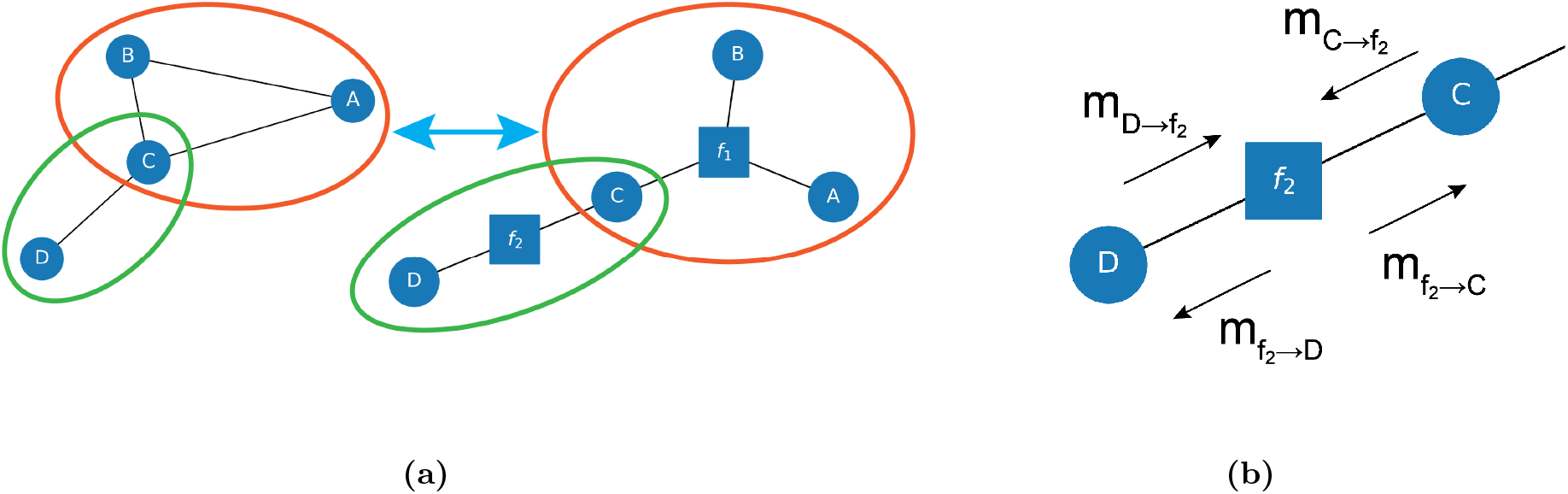
Depictions of clique factorization and message passing for an arbitrary set of variables where *A* - *D* represent variables and *f_i_* represent factors. (a) shows an MRF on the left with two cliques circled in orange and green converted into two factors circled in the same colors in the FG on the right. (b) shows all of the incoming and outgoing messages for factor *f*_2_ in the factor graph shown in (a).

This clique factorization can be shown more simply in another graph structure called a factor graph (FG). Factor graphs are bipartite graphs with nodes that are either factors or variables, and edges only connecting nodes between these two sets, but not within the sets (i.e. function node connected to variable node is the only allowed edge). This shows which variables are associated with which factor and how the factors are related through mutual connection to the same variable. This representation also simplifies the representation of the joint probability as shown in Eq. 5 because the product is now over factor nodes (*F*) and their associated edges (*X_a_*) instead of cliques (*C*(*G*)) and the nodes that compose them(*X_c_*).

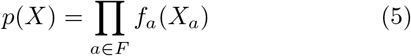

### II.C Belief Propagation

Thus far we have determined the relationship between our set of OPs using graphical lasso and converted this structure into a factor graph using clique factorization. We now know the OP sets we will use for each factor but have not learned the factors themselves or the joint probability they compose. We do so with the belief propagation (also known as sum-product) algorithm.^13,19,20^ This method updates an approximate factorized model *q*(*X*) of the joint distribution defined in Eq. 6 and composed of approximate factors 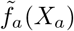:

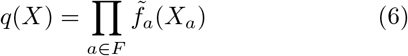

The objective of the belief propagation (BP) algorithm, first proposed by Pearl in 1982,^13^ is to approximate factors 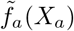 that will be iteratively updated so that their product *q*(*X*) approaches the true joint distribution *p*(*X*) in Eq. 5. To learn such factors, estimates of all other approximate factors are communicated to any single given factor at a time which is then adjusted depending on this “belief” about the other factors. The adjustment itself is carried out so that *p* and *q* mirror each other as closely as possible given the beliefs.

We now detail the BP equations that are used to calculate marginal probabilities and update messages. Here we index factors with *a, b* and variables with *i, j*. We denote sets of variables and factors with capital symbols *X* and *F* while the individual nodes comprising them are indicated with lowercase symbols *x* and *f* respectively. The sets *F* and *X* contain all nodes connected to the node with their index, for example *X_a_* represents all variable nodes *x* connected to the factor *f_a_*.

The central idea in our workflow is to mix kernel density estimation^21,22^ with traditional belief propagation. Specifically, we use kernel density estimation to model the marginal associated with each factor *p*(*X_a_*). We initialize all messages *m* to arrays of ones equivalent to passing no information about their factors. After this initialization step, our iteration begins as follows:

1. Select a factor to update, this is done at random for our simple case but could use more sophisticated methods.^23^
2. Apply Eq. 7 updating 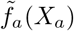 such that *q*(*X_a_*) = *p*(*X_a_*) from the kernel density estimator.

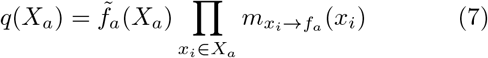
3. Apply Eq. 8 to update the factor’s outgoing messages.

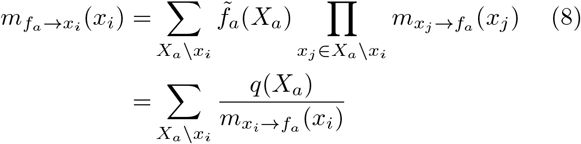
4. Apply Eq. 9 to update the messages outgoing from each variable associated with the factor.

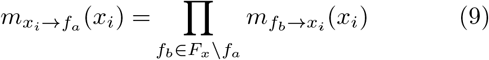
5. Repeat steps 1 – 4 until factors converge.

This procedure yields a factorized model for the joint probability that when marginalized will reproduce the marginal probabilities from kernel density estimation. This particular implementation of belief propagation was designed to be easily extended to use time evolving density estimators similar to that of metadynamics.^6,24,25^

## III RESULTS

In this section, we report detailed results of applying the methods described so far in this manuscript to two test biomolecules, namely capped Ala_3_ and Aib_9_.^14,16,26,27^ These respectively comprise 5 and 9 residues in explicit water, thus containing many complicated intertwined degrees of freedom. While the methods could easily be extended to consider solvent degrees of freedom as well,^28^ here we focus on the protein torsional degrees of freedom as the input OPs, numbering 6 and 18 for Ala_3_ and Aib_9_ respectively. For both systems we learn Markov random fields using graphical lasso. In addition, for Ala_3_ we learn a factor graph and use belief propagation to estimate the factors themselves. Thus we reduce the 6– and 18– dimensional probability distributions respectively to a combinations of pair and triplet interactions. The learned approximate lower-dimensional interactions are then analyzed and validated for both the systems in different ways, including intervention,^17^ enhanced sampling,^5,29^ and network analysis.

### III.A Capped Ala_3_

We performed 1 *μ*s long unbiased MD simulation (details in SI) recording three *ϕ* and three *ψ* dihedral angles every 200 fs. The trajectory of these six OPs was used to construct a MRF using graphical lasso as described in Sec. II A and this MRF was converted to a FG using clique factorization as described in Sec. II B. The MRF and the FG so learned are shown in Fig. 4.

**FIG. 4:**
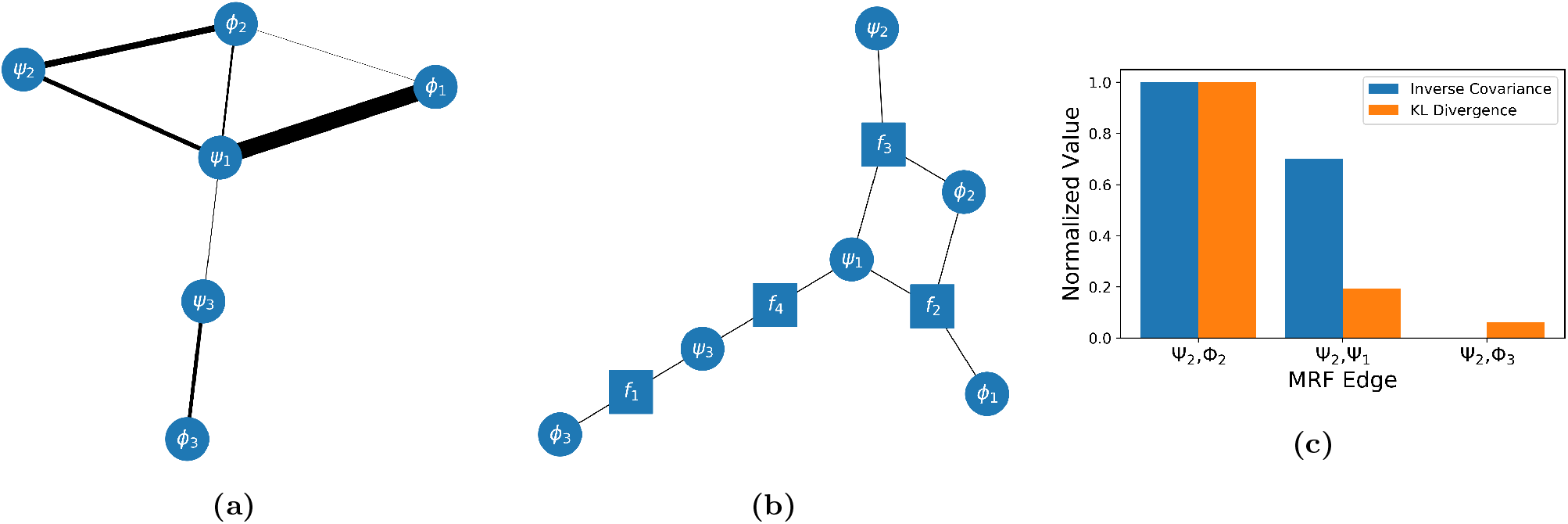
Graphical models of Ala_3_ calculated using a 1 *μ*s long molecular dynamics trajectory and comparison of the intervention results with the inverse covariance calculated with graphical lasso. (a) shows the MRF from graphical lasso with edge thickness showing the underlying inverse covariance, i.e. pairs of variables with thinner edges between them are more conditionally independent. (b) shows the FG calculated using clique factorization on (a). (c) shows the result of the intervention test that validates (a), by comparing the inverse covariance to the KL divergence between the observed probability during the intervention test and the independent probability model. We see that graphical lasso will favor drawing edges that clearly do not have the pairwise Markov property.

#### III.A.1 Validating the MRF through intervention

In order to validate this MRF, we perform a series of enhanced sampling simulations in the spirit of the intervention approach pioneered by Pearl.^17^ For each of these simulations we chose two OPs and restrained all other dihedrals in the MRF using a quadratic bias potential. This intervention procedure prevents any large fluctuations in the restrained OPs allowing us to directly assess the conditional independence hypothesis between the unrestrained OPs. Specifically, if a given pair of OPs are indeed conditionally independent given other OPs, then in the presence of such a restraint, the joint probability for this pair of OPs should be the same as the product of their marginal probabilities. Thus, in light of the MRF shown in Fig. 4, under respective constraints, we expect the joint probability to be closest to the product of constituent marginals for the pair Ψ_2_,Φ_2_, followed by Ψ_2_,Ψ_1_ and finally for Ψ_2_,Φ_3_. In order to assess this hypothesis, we use the Kullback–Leibler (KL) divergence^30^ to measure how close the sampled distribution is to the product of marginals. This represents an assessment of how well Eq. 1 models the sampled distribution. In Fig. 4 we show that indeed this is the case by comparing the KL divergence to the magnitude of the inverse covariance from graphical lasso. We see that the order of the inverse covariance and KL divergence match showing that graphical lasso did draw edges that lack conditional independence. The simulations were run for 1 *μ*s using all simulation details (except the restraint) identical to the initial Ala_3_ simulations, detailed in SI.

#### III.A.2 Validating the FG through enhanced sampling

With confidence in the MRF through the intervention procedure demonstrated in Fig. 4, these MRFs were then converted to factor graphs with up to triplet interactions. Given such a joint probability distribution expressed as product of terms up to triplets, we can directly use it to build a very high-dimensional bias potential that enhances the sampling simultaneously in all degrees of freedom, as defined below:

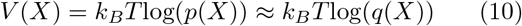

The central idea here is that the exact high-dimensional probability distribution *p*(*X*), if we knew it, would have led to the ideal bias for importance sampling which perfectly enhanced all fluctuations in the molecular system.^6^ The quality of the approximation *q*(*X*) in Eq. 6 can thus be ascertained by using it to construct an approximate bias as in Eq. 10. When added as a static bias to the system’s potential energy, a good approximation *p* ≈ *q* should then lead to spontaneous and enhanced back-and-forth movement between different metastable states. We convert the joint probability *q* to a bias potential using Eq. 10 with the caveat that we do not bias any regions of configurational space with a sampled probability of less that 0.001 in order to avoid adding bias to transition states. This threshold can be made higher or lower without affecting the nature of this discussion. The output of this calculation is a set of static external bias files to be used with PLUMED.^31–33^ The factor graph bias was used to conduct a 150 ns simulation using the same parameters used in the previous unbiased simulations for capped Ala_3_, implementing the bias through PLUMED.

As can be seen from the various time series in Fig. 5, the simulations conducted using the static bias potential enhanced the fluctuations of all individual OPs. We also consider the transition rates for all of the 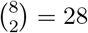 transitions between the eight dominant metastable states, corresponding to the positive and negative regions of the Ramachandran plot for each of the three Alanine residues. These are shown in Fig. 5(g) as the average interconversion or commute time between different states.^26^ When compared to the unbiased sampling, the factor graph biased simulations have significantly enhanced sampling (at least twice as fast, and up to 350 times faster among those sampled in both simulations) along 22 of the 28 transitions. The only transitions that are not enhanced are the ones that were fast to begin with except the notable exception of *S*_2_ to *S*_5_ which was not sampled in 150 ns of simulation with the factor graph bias. Interestingly, the low barrier transitions are not significantly slowed down due to biasing, which has been a concern in enhanced sampling.^34^ Finally, in addition to the bias associated with the factor graph, we repeated both the bias generation and biased simulation for a factor graph without any edges which corresponds to biasing each OP in parallel without considering any correlations. These simulations are used for additional comparison in the SI.

**FIG. 5:**
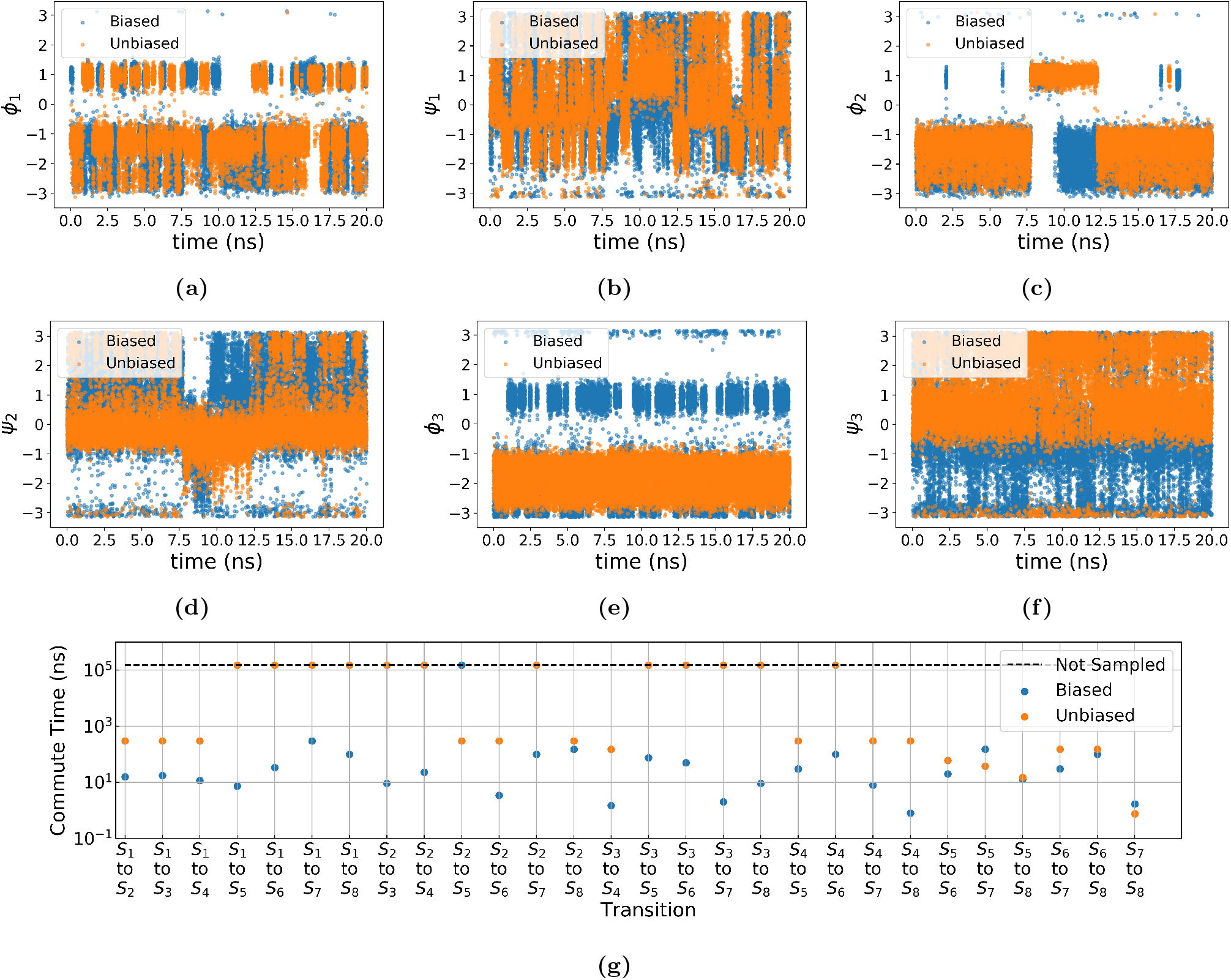
(a)–(f) Trajectories of six OPs for Ala_3_ for factor graph biased (blue) and unbiased (orange) simulations. It is clearly evident that the factor graph based biased simulations achieve spontaneous back-and-forth movement across high-dimensional configuration space. The acceleration relative to unbiased MD is quantified in (g), which shows the commute time for pathways between every pair of states for factor graph biased (blue) and unbiased (orange) simulations of capped Ala_3_. We use the state definitions described in Ref. 26. Points on the dotted line were not sampled and do not correspond to a commute time of 10^5^.

**FIG. 6:**
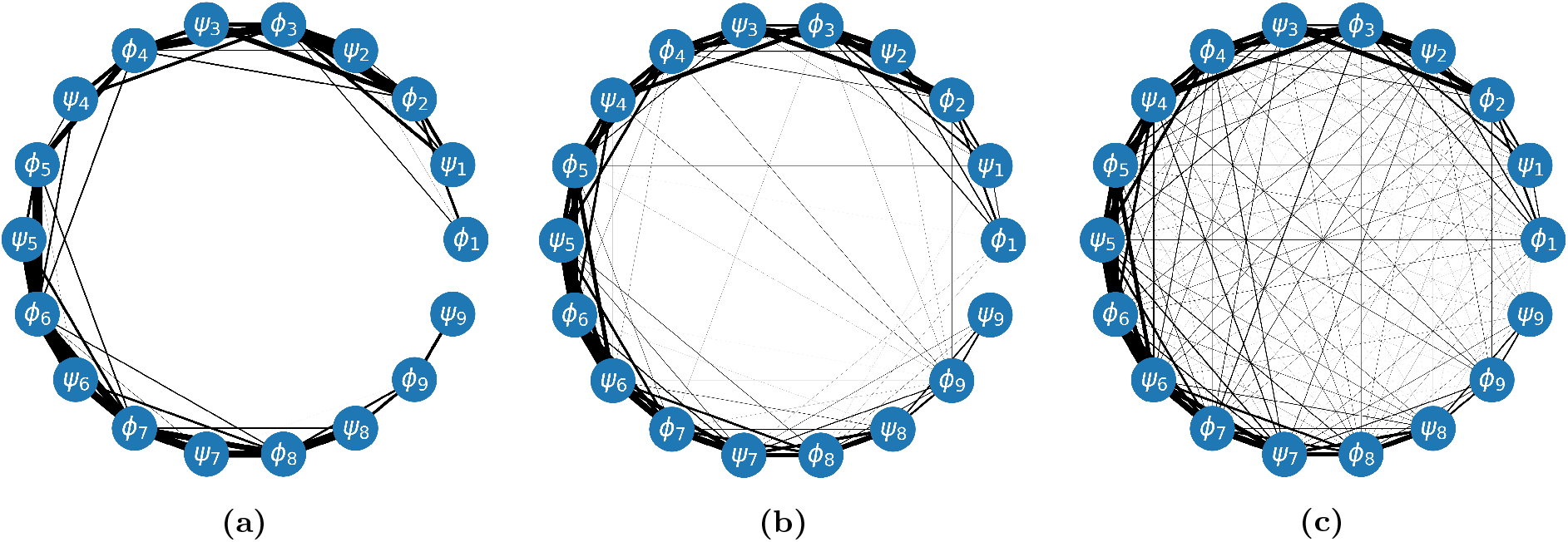
Markov random fields calculated using trajectories of Aib_9_’s dihedral angles with edge thickness showing the magnitude of the inverse covariance. (a) corresponds to a penalty parameter lower than that obtained with 5-fold cross validation. (b) corresponds to the cross-validated penalty parameter value. (c) corresponds to a penalty parameter lower than the cross-validated value.

### III.B Aib_9_ Peptide

Aib_9_ is a *α* aminoisobutyric acid-based 3_10_-helix^15,16,27^ that displays conformational transitions at different timescales ranging from picoseconds to microseconds, and that may very well vary with solvent. Here we study Aib_9_ in explicit water with 200 ns of simulation (see SI for details).^35^ The most distinct of these is the transition between right- and left-handed helices, as shown in Fig. 1. Aib_9_ requires nine *ϕ* and nine *ψ* dihedrals for a complete description of the conformational dynamics. The trajectory of these 18 OPs was used to calculate Markov random fields shown in Fig. 6 which shows the change in graph structure as the graphical lasso penalty parameter is tuned.

The secondary structure of Aib_9_ causes these OPs to be correlated in larger groups which are not amenable to our procedure limiting the factor graph to at most triplet interactions. This means that additional techniques^36–39^ will be required to reduce the dimensionality of the OPs associated with larger factors in more complex systems which will be the subject of future investigation. Nevertheless, we can infer interesting conclusions regarding the underlying network governing the conformational dynamics of Aib_9_ from Fig. 6. Firstly, irrespective of the value of the penalty parameter *λ* in Eq. 3, we never find the two ends of the peptide to be connected with a significant inverse covariance, in contrast to the MRF we found for the smaller capped Ala_3_ peptide. Secondly, the interior residues of the peptide display thicker edges than the ones on the outside, demonstrating that inner residues move in a more correlated and conditionally dependent manner. Thirdly, the MRF is roughly symmetric as one approaches the center from either end i.e. clockwise from Ψ_9_ or anticlockwise from Φ_1_. This is indeed in agreement with measurements from sophisticated enhanced sampling and much longer (i.e. at least several microsecond) simulations, which suggest that Aib_9_ is achiral and both the left-hand right-handed helices have same free energies.^16,27^ We also find that applying the Girvan-Newman algorithm^40^ to the sparse and cross-validated MRFs yields two communities which are split by the center of the protein. These communities suggest the edges crossing the two sides of the protein are involved in many shortest paths between nodes and in turn the dynamics of the central residues are crucial for conformational change. It is interesting to note that MRF trained on the first 80 ns or longer gives the same result as the trajectory has sufficiently sampled both right- and left-handed helices.

## IV DISCUSSION

In this work, we have developed and applied a probabilistic graphical models based framework for making sense of high-dimensional data arising in molecular simulations. This makes it possible to learn reliable approximations to high-dimensional molecular distributions in a tractable manner. Our approximations here are restricted up to a family of single, pair and triplet interactions but could be easily extended further if needed.

We validated this framework through detailed analysis of two biomolecular tests systems. First, we used it to learn a factor graph structure for the small peptide capped Ala_3_ that captures inter-residue correlations relevant to various conformational transitions in this molecule. Inspired by recent methods^41,42^ which conduct enhanced sampling biasing multiple independent components, we demonstrate how our framework can be used to divide a large number of order parameters (OPs) into groups for biasing. As such, we use our factor graph structure along with belief propagation to construct a static bias for enhanced sampling. Using this bias, with no time dependence, we were able to enhance the fluctuations along each OP as well as accelerate the sampling of almost all state-to-state transitions with spontaneous back-and-forth movement between states. While this static bias approach is arguably not a practical enhanced sampling method, it illustrates the importance of including correlations in the biasing variables as well as the possibility to use factorized models with more sophisticated enhanced sampling methods.^39,41^ For this system we also demonstrate an intervention procedure^17^ that directly tests the predictions of inter-residue correlations learned in our framework. As a second illustrative system we also model the peptide Aib_9_ using a Markov field constructed with graphical lasso. This allows us to highlight the interpretability of probabilistic graphical models applied to molecular simulations of this peptide as it undergoes an interesting left-handed to right-handed helix transition. This graph structure highlights features of Aib_9_ such as the the difference in flexibility between central and outer residues, and the symmetry of correlations between residues.

We conclude by noting that despite the limitations of our simple illustrative strategies, this work can also be combined with sampling methods that are used to determine effective sets of basis functions for describing molecular systems^43^ or those that are used for dimensionality reduction.^36,39,44^ We envision that these methods will be used to develop an automated framework for sampling biomolecules that takes a starting structure and runs enhanced sampling simulations without the necessity for expert knowledge. This combination should allow simulation of new systems where traditional low-dimensional biasing variables are unsuccessful in a high-throughput manner, for instance biomolecular complexes with more than one constituent. Finally, we highlight that the framework developed in this manuscript is also closely related to the topic of causal inference.^45,46^ The graphs learned for these molecular systems can be converted into directed Bayesian networks when combined with information about whether past states of an order parameter are predictive of future states of another. This would further improve the ability to understand molecular systems as it allows us to make not just correlational but causal and mechanistic claims.

## Notes

The code needed to reproduce the models used in this work will be made available at github.com/tiwarylab. The input files necessary to reproduce the simulations done in this work will be made available on PLUMED NEST.^47^

## ACKNOWLEDGMENTS

This work was supported by the National Science Foundation, Grant No. CHE-2044165. ZS was also supported by University of Maryland COMBINE program NSF award DGE-1632976. This work used XSEDE Bridges through allocation TG-CHE180053, which is supported by National Science Foundation grant number ACI-1548562. We also thank UMD’s Deepthought2 and MARCC’s Bluecrab HPC clusters for computing resources. We would like to thank Shams Mehdi for the Aib_9_ trajectory. We would also like to thank Pavan Ravindra, Yihang Wang, Michael Strobel, and Nathan Zimmerberg for discussion and feedback about the direction of this project.

## Notes

### Competing Interest Statement

The authors have declared no competing interest.

