## Supplemental Information for "Making high-dimensional molecular distribution functions tractable through Belief Propagation on Factor Graphs"

**Overview**

This document provides details of simulations used for both the systems in the manuscripts, as well as further explanatory figures numbered Fig. S1 to S3.

### A. Initial Simulations

In order to calculate graph structures for a specific system and associated set of variables we must first obtain simulation trajectories to analyze. All molecular dynamics simulations were run through the GROMACS package patched with PLUMED version 2.4.2[1,2] with a 2 fs time step. All order parameters (OPs) for both systems were recorded every 200 fs. Capped $Ala_3$[3] ($Ace - Ala_3 - Nme$) was simulated for 1 $\mu$s using forcefield parameters determined in Ref. 3. Temperature was kept at 400 K using the velocity rescale thermostat.[4] The nonbonded interactions, including electrostatics, were calculated with a 10 Å cutoff.

$Aib_9$ peptide[5] ($H_3C - CO - (NH - C_\alpha(CH_3)_2 - CO)_9 - CH_3$) simulations were run for 200 ns using forcefield parameters determined in Ref. 6 using the charmm36 forcefield.[7] Temperature was kept at 500 K using the Nose-Hoover thermostat.[8] Pressure was kept at 1 bar using the Parrinello-Rahman barostat.[9] The nonbonded interactions were calculated using a Verlet cutoff scheme with a 12 Å cutoff.[10]

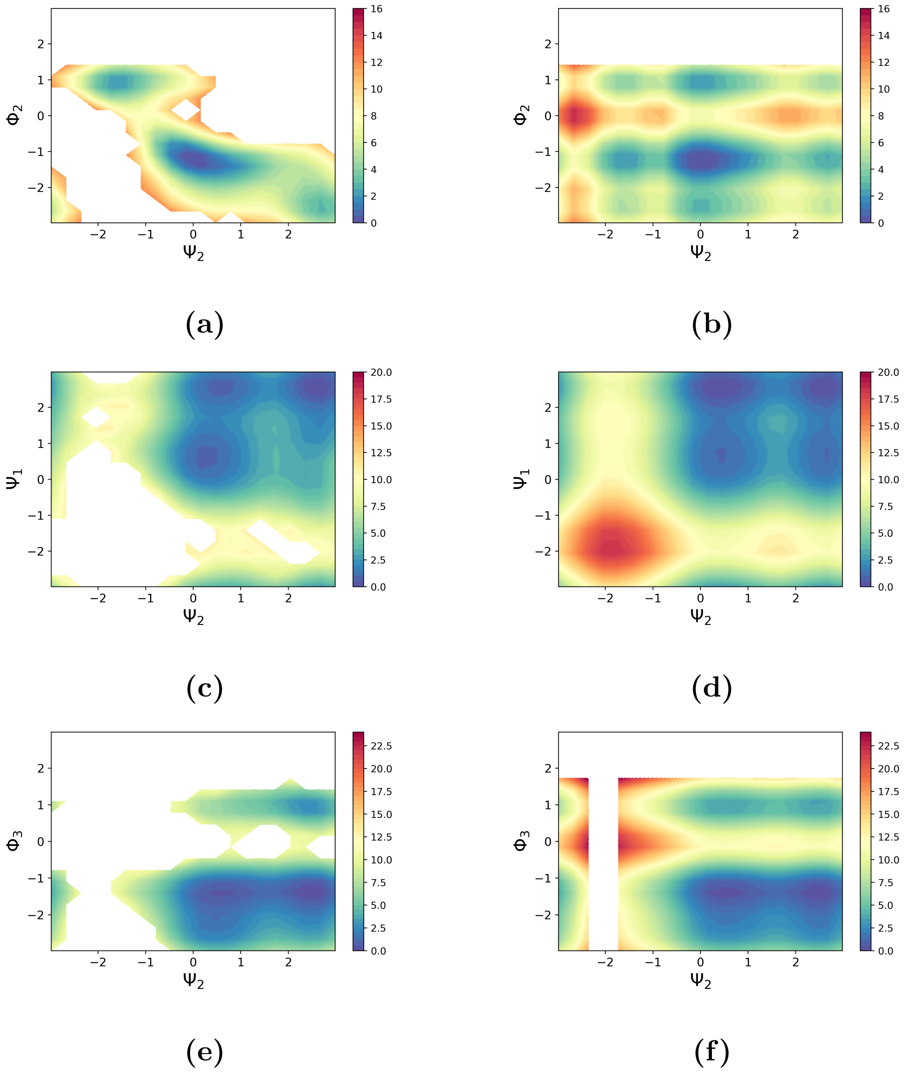


**FIG. S1:** Comparison of simulated free energy surfaces ((a), (c), and (e)) to those expected if each OP was independent ((b), (d), and (f)) with contour levels shown every .5 $k_BT$. The independent free energies were obtained by multiplying the marginal probabilities for both of the OPs in a given plot.

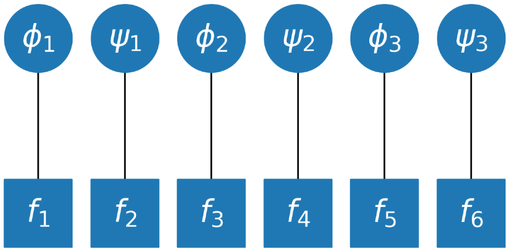


**FIG. S2:** Factor graph corresponding to biasing each variable independently in parallel. Used to run an additional enhanced sampling simulation for comparison with the original factor graph biased simulation.

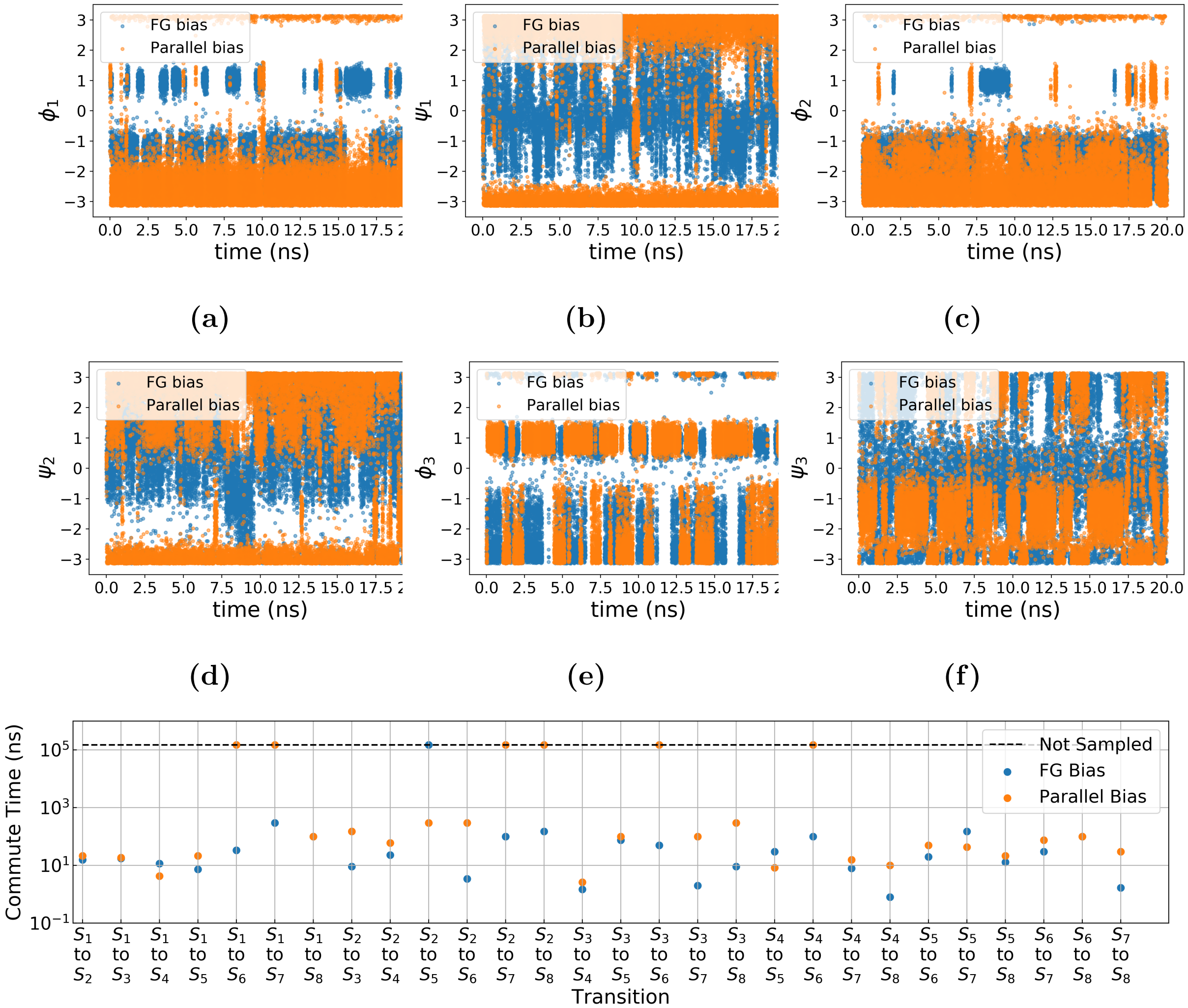


**FIG. S3:** (a)–(f) Trajectories of six OPs for $Ala_3$ for factor graph biased (blue) and parallel biased (orange, using the FG in Fig. S2) simulations. The acceleration relative to parallel bias is quantified in (g), which shows the commute time for pathways between every pair of states for factor graph biased (blue) and unbiased (orange) simulations of capped $Ala_3$. We use the state definitions described in Ref. 11. Points on the dotted line were not sampled and do not correspond to a commute time of $10^5$. This figure corresponds to Fig. 5 in the main text comparing the factor graph bias with parallel bias simulations instead of unbiased.
